# Personalized genome assembly for accurate cancer somatic mutation discovery using cancer-normal paired reference samples

**DOI:** 10.1101/2021.04.09.438252

**Authors:** Chunlin Xiao, Zhong Chen, Wanqiu Chen, Cory Padilla, Li-Tai Fang, Tiantian Liu, Valerie Schneider, Charles Wang, Wenming Xiao

**Affiliations:** National Center for Biotechnology Information, National Library of Medicine, National Institutes of Health, 45 Center Drive, Bethesda, MD 20894; Center for Genomics, Loma Linda University School of Medicine, 11021 Campus St., Loma Linda, CA 92350; Dovetail Genomics, 100 Enterprise Way, Scotts Valley, CA 95066; Bioinformatics Research & Early Development, Roche Sequencing Solutions Inc., 1301 Shoreway Road, Belmont, CA 94002; The Center for Drug Evaluation and Research, U.S. Food and Drug Administration, Silver Spring, MD, USA

## Abstract

The use of personalized genome assembly as a reference for detecting the full spectrum of somatic events from cancers has long been advocated but never been systematically investigated. Here we address the critical need of assessing the accuracy of somatic mutation detection using personalized genome assembly versus the standard human reference assembly (i.e. GRCh38). We first obtained massive whole genome sequencing data using multiple sequencing technologies, and then performed *de novo* assembly of the first tumor-normal paired genomes, both nuclear and mitochondrial, derived from the same donor with triple negative breast cancer. Compared to standard human reference assembly, the haplotype phased chromosomal-scale personalized genome was best demonstrated with individual specific haplotypes for some complex regions and medical relevant genes. We then used this well-assembled personalized genome as a reference for read mapping and somatic variant discovery. We showed that the personalized genome assembly results in better alignments of sequencing reads and more accurate somatic mutation calls. Direct comparison of mitochondrial genomes led to discovery of unreported nonsynonymous somatic mutations. Our findings provided a unique resource and proved the necessity of personalized genome assembly as a reference in improving somatic mutation detection at personal genome level not only for breast cancer reference samples, but also potentially for other cancers.

## Introduction

Accurately detecting somatic mutations and subsequently understanding genomic instability in cancer are critical for precision cancer therapies, and many genomics studies, including tremendous efforts from the renowned TCGA and ICGC consortia, have been directed to investigate genomic instability of cancer, and greatly improved our understanding of cancer biology [1–4]. Most recently, tumor-normal pair reference samples and reference callsets were established by the SEQC-II consortium for benchmarking somatic mutation detections from different sequencing platforms and bioinformatic analysis methods [46–47]. This standard is an indispensable foundation and resource for assessing the accuracy and reproducibility of somatic mutation profiling in cancer cell lines.

To date, discovering cancer somatic events and defining high-confidence reference somatic callsets mainly rely on a standard human reference assembly (such as GRCh38) as a benchmark for sequencing analysis. However, the GRCh38 has its own limitations. Although of very high quality, it remains incomplete due to some unresolved assembly issues and persistent gaps, including those at centromeres, telomeres, and heterochromatic regions [5–8]. Incorrect or missing sequences in GRCh38 reference assembly may lead to failed or spurious read mapping and unreliable subsequent analysis results (namely reference bias [9]). Moreover, the human reference assembly was derived from the DNAs sourced from multiple individuals, though approximately 70% of the GRCh38 sequences were contributed by a single Africa-European admixed male (RP11) [6]. Such mosaic haplotype representation in the reference assembly may complicate the identification of somatic variations from cancer samples; therefore, use of a *de novo* assembly of the personalized genome, rather than the standard reference assembly, for confident cancer mutation discovery has been advocated [5, 7, 10]. There is more to be learned from direct comparison of the tumor genome to the normal genome from which it is derived, than to an unrelated, random, mosaic genome like the latest GRCh38 reference genome.

The advancements of DNA sequencing technologies provide an extraordinary opportunity to perform *de novo* assembly for individual genomes at affordable cost. Particularly, the breakthrough of long range DNA sequencing from the third-generation sequencing technologies now makes it possible to accurately assemble individual genomes to near completion, which has been done for several samples, including HX1 [11], AK1 [12], NA12878 [13], CHM13 [14], and HG002 [15]. These studies demonstrated the recent advancements in genome assembly methods and subsequent germline variant detection. However, using personalized genome as reference for somatic mutation detection in cancer, particularly in cancer-normal paired samples, has not been systematically investigated. Here we present our work that combined multiple sequencing technologies, including Illumina short reads, 10X Genomics linked reads, PacBio long reads, and Hi-C (high-throughput chromosome conformation capture) reads, to reconstruct, to our knowledge, the first tumor-normal paired genomes [7]. Derived from the same individual, HCC1395 cancer cell line, and a matched B lymphocyte cell line HCC1395BL [16] as well-studied reference samples by FDA-led SEQC-II consortium [46–47], represent one of the most important tumor-normal models for triple negative breast cancers (TNBC). This work will enable us to comprehensively assess the performance of somatic mutation detection with respect to the underlined reference assemblies being used, as well as pinpoint the complete spectrum of genomic alterations, using a personal genome as the reference.

## Results

### Overall study design and establishing a reference-grade personal genome assembly

We used five platforms for sequencing the normal reference sample (HCC1395BL B Lymphocyte cell line) and three platforms for sequencing the tumor reference sample (HCC1395 breast cancer cell line) from the same donor (Figure 1, Top). Using data from multiple sequencing technologies, including short reads, linked reads, and long reads (Supplementary Table 1), we built a workflow for establishing a *de novo* assembled genome (Figure 1, Bottom-Left). First, we generated initial assemblies for the HCC1395BL, using canu [17] for PacBio long reads and using Supernova [18] for 10X Genomics linked reads, respectively. Supernova assemblies contained many smaller contigs (size less than 10 kb) than PacBio canu assemblies (Supplementary Table 2). Although the N50s and the largest scaffold of the Supernova assembly were much higher than the PacBio contig assembly (Supplementary Figure 1, Supplementary Figure 2), the latter was more complete as measured via Benchmarking Universal Single-Copy Orthologue (BUSCO) genes (Supplementary Figure 3). Additionally, a greater number of complete RefSeq protein-coding genes mapped to the PacBio assembly, and more base pairs from this assembly could be mapped to the GRCh38 reference (Supplementary Table 2). Taken together, these indicated that the overall quality of PacBio canu assembly was higher than Supernova assembly, particularly when gene content and the completeness are the primary concerns.

**Table 1.** Summary of quality assessments of assemblies from different scaffolding strategies.

|  | # scaffolds | # bps from scaffolds | Top50 | N50 | L50 | Largest scaffold length (bps) | # scaffolds on GRCh38 | # novel scaffolds | # bps from novel scaffolds | # Complete BUSCO (4,104 BUSCO) | # NMs 95+% mapped (50,052 NMs) | # NRs 95+% mapped (15,544 NRs) |
| --- | --- | --- | --- | --- | --- | --- | --- | --- | --- | --- | --- | --- |
| PB_canu (contigs) | 2,828 | 2,904,842,414 | 1,356,447,278 (46.69%) | 13,480,407 | 57 | 62,208,403 | 2,526 | 302 | 16,403,702 | 3,890 | 49,287 | 15,115 |
| PB_canu + 10X_arcs (scaffolds) | 2,032 | 2,904,931,213 | 2,067,627,443 (71.17%) | 35,058,531 | 26 | 121,623,092 | 1,764 | 268 | 14,867,391 | 3,800 | 49,570 | 15,207 |
| PB_canu + HiC_salsa (scaffolds) | 1,891 | 2,905,381,691 | 2,377,981,926 (81.84%) | 46,871,224 | 19 | 180,772,639 | 1,617 | 274 | 15,303,825 | 3,892 | 49,495 | 15,177 |
| PB_canu + 10X_arcs + HiC_salsa (scaffolds, HCC1395BL_v1.0) | 1,645 | 2,905,196,510 | 2,691,295,119 (92.62%) | 69,970,292 | 14 | 181,209,810 | 1,406 | 244 | 14,104,388 | 3,889 | 49,613 | 15,227 |

**Figure 1.**
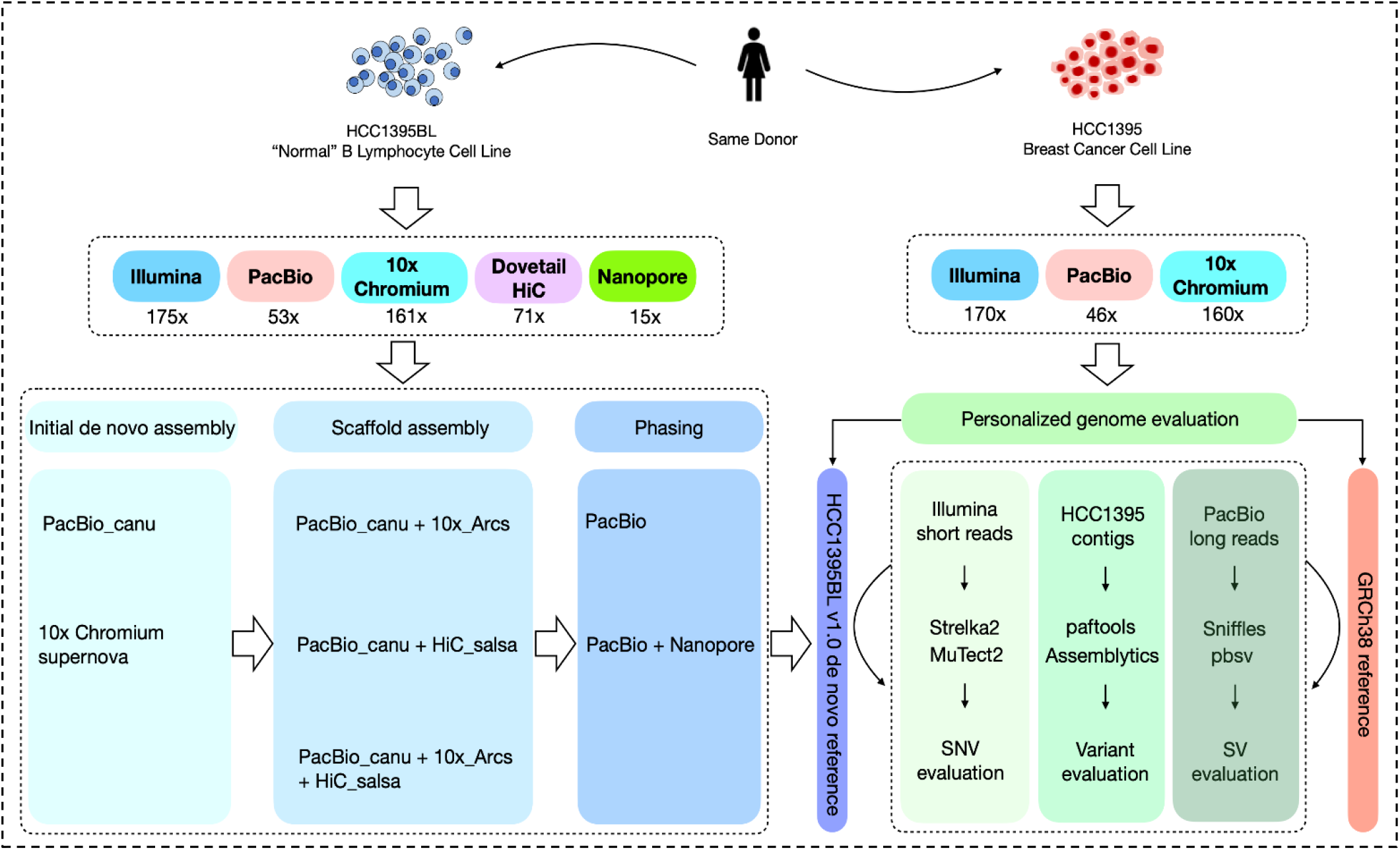
Schematic diagram of study design, DNA sequencing of the tumor-normal paired reference cell lines (HCC1395BL and HCC1395), *de novo* genome assembly of the cell lines, and assessment of somatic mutation detection using standard GRCh38 genome and personal genome assembly as references, respectively.

We then selected the PacBio contig assembly of HCC1395BL for further scaffolding. In general, two steps of scaffolding, first using 10X Genomics linked reads with ARCS, followed by Hi-C reads with SALSA (PacBio_canu + ARCS + SALSA), produced a better scaffolded assembly than using one-step scaffolding only (either PacBio_canu + ARCS or PacBio_canu + SALSA). The final scaffold assembly (hereafter referred to as HCC1395BL_v1.0) was the one with the highest Top50 (see “Methods”) and scaffold N50 values, the largest scaffold size, and the greatest numbers of mapped complete BUSCOs and RefSeq transcripts (Table 1). HCC1395BL_v1.0 consisted of 1,645 scaffolds totaling 2.9 Gb, of which 2.69 Gb (92.75%) is from Top50 scaffolds, with a scaffold N50 size of 69.97 Mb, in comparison to scaffold N50s 67.79 Mb for GRCh38 [6], and 44.84 Mb for AK1, a recent assembly from a diploid sample [12], respectively.

Both HCC1395BL_v1.0 and AK1 had a very similar assembly size (2.90 Gb), but HCC1395BL_v1.0 had fewer scaffolds (1,645 vs. 2,832), a smaller L50 (14 vs. 21), and had much greater Top50 (2.69 Gb vs. 2.26 Gb), N50 (69.97 Mb vs. 44.84 Mb), and largest scaffold size (181.21Mb vs. 113.92 Mb). Moreover, the HCC1395BL_v1.0 assembly contained more complete RefSeq NM (protein-coding) transcripts (49,613 vs. 49,432) and RefSeq NR (non-protein-coding) transcripts (15,227 vs. 15,089).

Consistency analysis (see “Methods”) with the GRCh38 primary assembly (alt_loci excluded) showed that five chromosomes (chr4, chr8, chr14, chr18, and chr20) were near-completely covered by single scaffolds. The largest HCC1395BL_v1.0 scaffold (Scaffold_1 181.21Mb) covered more than 95% of GRCh38 chromosome 4. Four chromosomes (chr2, chr3, chr12, and chr19) were just broken at centromere regions. Several other chromosomal arms (chr1p, chr5p, chr6q, chr9p, chr10p, chr21q, and chrXq) were also covered by single HCC1395BL_v1.0 scaffolds (Figure 2).

**Figure 2.**
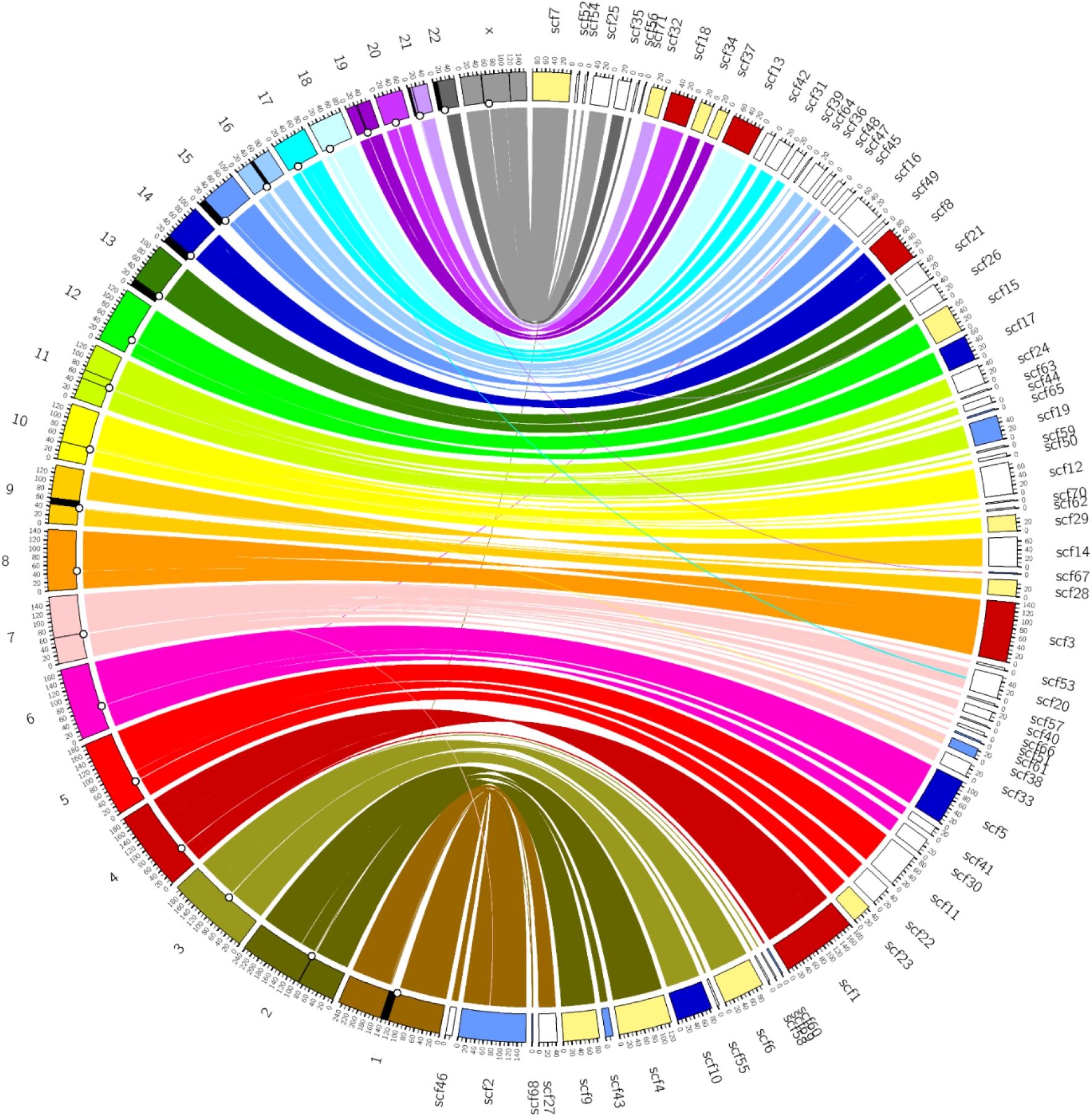
A Circos consistency plot of HCC1395BL_v1.0 (right side) against GRCh38 reference genome (left side). Included were 71 largest scaffolds with 2+ Mbp that accounted for 2,775,074,314 bps (95.51%). Shown here were alignments with coverage of 100Kb+ and mapping quality 60 on GRCh38 using minimap2. Centromeres were marked with black circle on the inner circle of GRCh38 chromosomes. Black regions on the chromosomes were GRCh38 gaps with 100Kb+ in size. Five chromosomes were near-completely covered by single scaffolds (Scaffold_1 for chr4, Scaffold_3 for chr8, Scaffold_8 for chr14, Scaffold_13 for chr18, and Scaffold_18 for chr20, were colored with red color). Four chromosomes (chr2, chr3, chr12, and chr19) were broken only at centromere regions (near-completely covered by just two scaffolds). Centromere-crossing scaffolds were colored with light blue. Scaffolds (Scaffold_5, Scaffold_10, and Scaffold_17) covering one arm and crossing the centromere were colored dark blue. Scaffolds with near full coverage of one arm were covered with yellow.

Phasing analysis showed that 3.13 out of 3.17 million heterozygous sites were considered phased, and 6,368 phased blocks accounted for 2.42 Gb of HCC1395BL_v1.0, with the longest phased block 6.37 Mb (Supplementary Table 3). Approximately 15-fold coverage of Nanopore long reads were used to further extend phasing to 2.54 Gb. The total number of phased blocks was subsequently decreased to 3,204 from 6,368 blocks, and the longest phased block was greatly improved, increasing from 6.37 Mb to 20.45 Mb (Supplementary Table 3). With the phased assemblies (haplotype1 and haplotype2) for HCC1395BL cell line, we were able to call 4,115,622 germline SNVs in diploid regions of autosomal chromosomes.

As a comparison, we also generated a *de novo* assembly for the HCC1395 cancer cell line, using both canu for PacBio long reads and Supernova for 10X Genomics’ linked reads. The resulting HCC1395 assembly was more fragmented than HCC1395BL assembly (Supplementary Table 2), mainly due to high level of chromosomal aneuploidy and structural variations in this cancer cell line [19].

### Personal genome assembly includes individual specific haplotypes for clinically relevant genes, and eliminates special handling of alt_loci/patches

We aligned a NCBI RefSeq transcript set (excluding all pseudogenes and genes from chromosome Y) to HCC1395BL_v1.0. 19,303 of 19,325 (99.89%) RefSeq protein-coding genes could be successfully mapped onto HCC1395BL_v1.0 with minimum 95% alignment identity and 50% alignment coverage, while 19,164 of 19,303 (99.27%) these genes aligned at least 95% coverage (Figure 3A and Supplementary Table 4). Among RefSeq non-protein-coding genes, 10,049 of 10,061 (99.88%) genes could be aligned to HCC1395BL_v1.0 successfully with minimum 95% identity and 50% coverage, with 9,958 of 10,049 (99.09%) those genes covered at more than 95% in length (Figure 3A and Supplementary Table 4).

**Figure 3.**
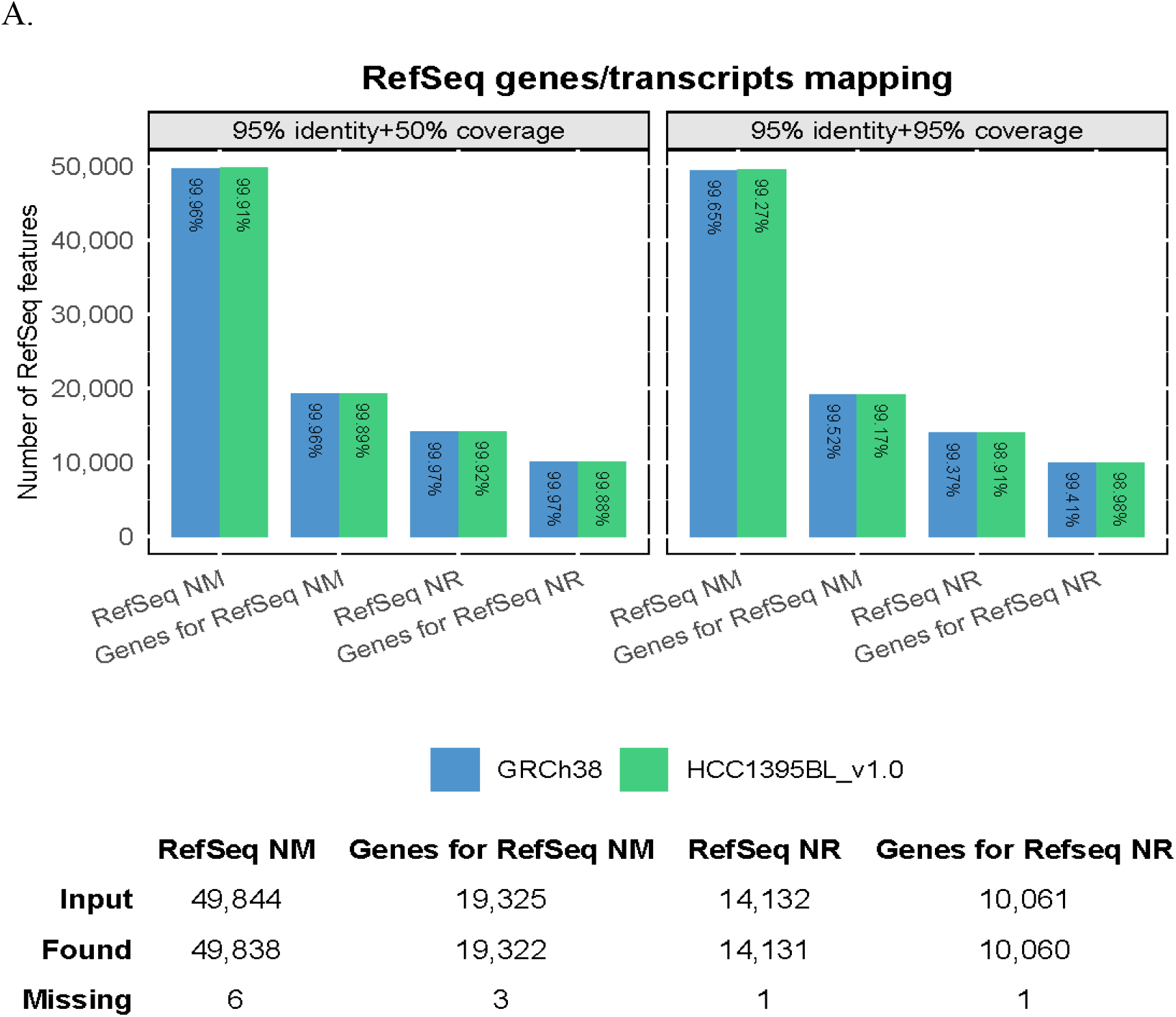
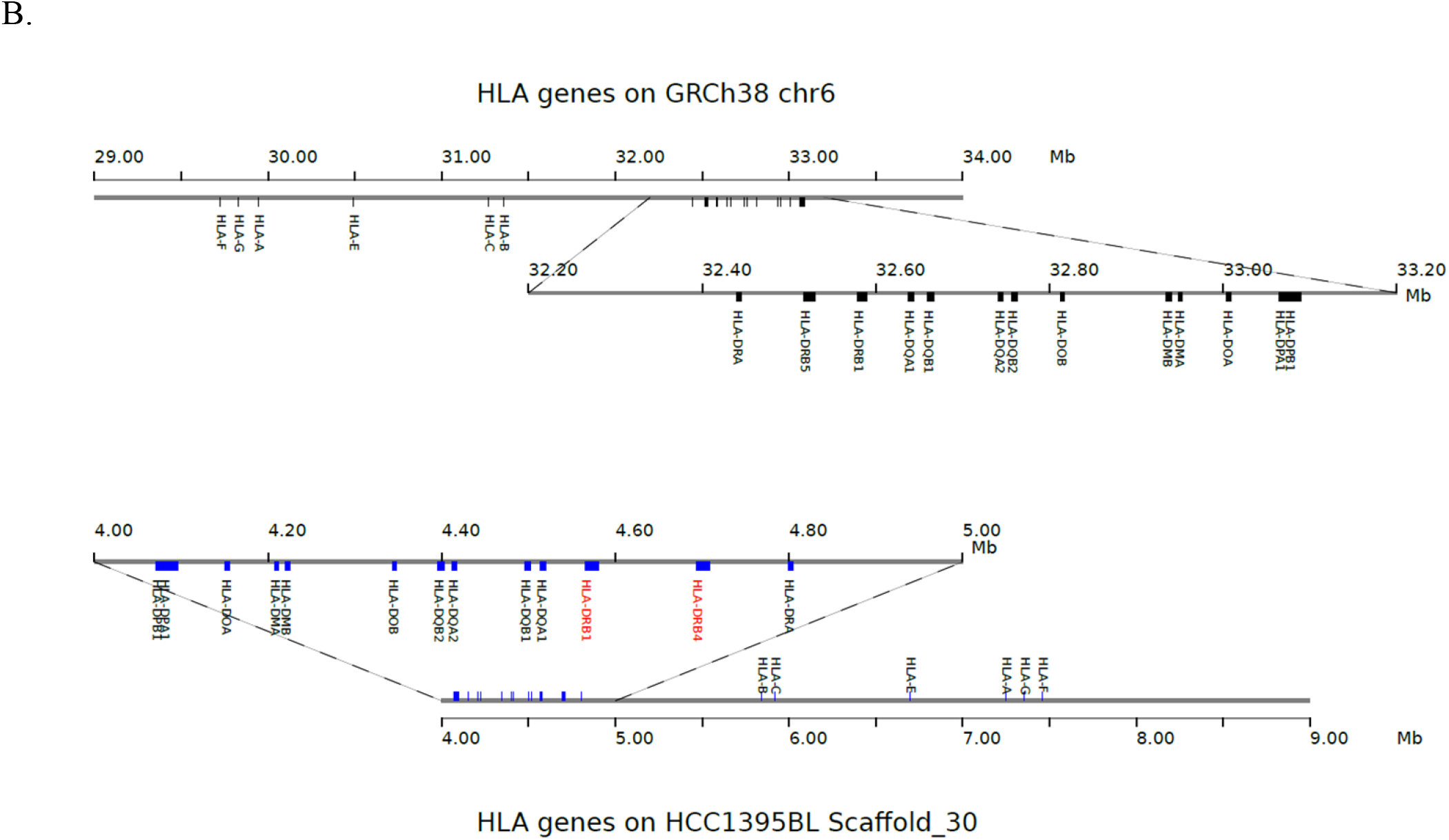
(A) The summary of RefSeq genes/transcripts mapping on HCC1395BL_v1.0 and GRCh38; (B) HLA genes on Scaffold_30 of HCC1395BL_v1.0 in comparison to those on chromosome 6 of GRCh38 primary assembly. The haplotype of HLA-DRB (labels in red) in HCC1395BL_v1.0 consists of HLA-DRB1 and HLA-DRB4 genes (human HLA-DR53 haplotype group), while GRC38 primary assembly contains HLA-DRB1 and HLA-DRB5 genes (human HLA-DR51 haplotype group). The human HLA-DR53 haplotype was represented only in GRCh38 ALT_REF_LOCI sequences. Scaffold_30 was reverse complement mapped on GRCh38, but the order of HLA genes on chromosome or scaffold was identical between GRCh38 and HCC1395BL_v1.0.

We next compared HLA gene family coverage in GRCh38 and HCC1395BL_v1.0, as an individual could have a very different HLA haplotype from another individual or the standard reference genome like GRCh38. HLA genes are located on the 6p region of chromosome 6, and previous cytogenetic analysis showed that this region from the HCC1395BL cell line was essentially haploid [46–47]. From the RefSeq gene set, 19 HLA genes (25 coding transcripts) are annotated on chromosome 6 of the GRCh38 primary assembly. We successfully identified all the HLA genes and corresponding transcripts in HCC1395BL_v1.0 and found that they are located on a single scaffold (Scaffold_30) aligning at minimum identity 95% and the minimum alignment coverage 95%. Notably, the order of the HLA genes on this scaffold is identical to that on GRCh38 (Figure 3B). The only difference is the haplotype of HLA-DRB genes between HLA-DRA and HLA-DQA1, as the region of HLA-DRB genes is extremely divergent [20]. The haplotype of HLA-DRB in GRCh38 primary assembly was represented by HLA-DRB1 and HLA-DRB5 genes (human HLA-DR51 haplotype group [21]), but the HLA-DRB haplotype in HCC1395BL_v1.0 consists of HLA-DRB1 and HLA-DRB4 genes (human HLA-DR53 haplotype group), which is similar to the HLA-DRB haplotypes represented on GRCh38.p13 ALT_REF_LOCI_4 (NT_167246.2) and GRCh38.p13 ALT_REF_LOCI_7 (NT_167249.2). This demonstrated that the *de novo* assembly and scaffolding of HCC1395BL_v1.0 performed well on such hypervariable/complex regions that harbors HLA genes.

We also evaluated other clinically relevant genes whose only representations in GRCh38 are on alternate locus scaffolds, included in the reference to capture population diversity. HCC1395BL_v1.0 included GSTT1 (Glutathione S-transferase theta 1), and KIR2DL5A (Killer cell immunoglobulin like receptor, two Ig domains and long cytoplasmic tail 5A), two genes that are not included in the haplotypes represented on the chromosomes of the GRCh38 primary assembly. GSTT1, a gene previously localized to chromosome 22 of GRCh37 primary assembly, is found on alternate locus scaffold (NT_187633.1) in GRCh38. Likewise, for KIR2DL5 gene, the haplotype represented on Chromosome 19 with un-localized genomic contig (NT_113949.1) on the GRCh37 primary assembly included KIR2DL5A, but in the GRCh38 assembly, it is only found in alt_loci and novel patches (NT_113949.2). The changes in representations of these genes from GRCh37 to GRCh38 (or the future version) present analysis challenges when switching between versions of the standard reference genome. Additionally, because these genes are represented only on alt_loci and patches in GRCh38, and most existing tool chains do not handle those alternate locus scaffolds, they are consequently more difficult to study, their exclusion from analysis presents a heightened risk for misinterpretation of results. In contrast, as the individual specific haplotypes for these clinically relevant genes were correctly represented in the personalized assembly, no special handling of alt_loci or patches would be needed to assess them if HCC1395BL_v1.0 is used as reference as opposed to GRCh38.

### Personal genome assembly as reference enables better read mapping and accurate cancer somatic mutation detection

We then used HCC1395BL_v1.0 as a reference for Illumina short read mapping and somatic variant discovery (Figure 1, Bottom-Right). While the mapping rates of short reads to the GRCh38 primary assembly (alt_loci excluded) and HCC1395BL_v1.0 were similar, we observed overall improved read mapping on HCC1395BL_v1.0 as opposed to GRCh38 (Figure 4A and Supplementary Table 5). For instance, a slightly higher percentages of properly-paired reads being mapped for both normal and tumor samples were seen on HCC1395BL_v1.0. Notably, the numbers of the improperly paired reads were reduced by 33~36%, whereas the numbers of the mismatches for the mapped reads were decreased by about 16%. In addition, the numbers of read alignments with soft-clipping were down by 12~13%, while the numbers of read alignments with hard-clipping were down by 28~31% (Supplementary Table 5).

**Figure 4.**
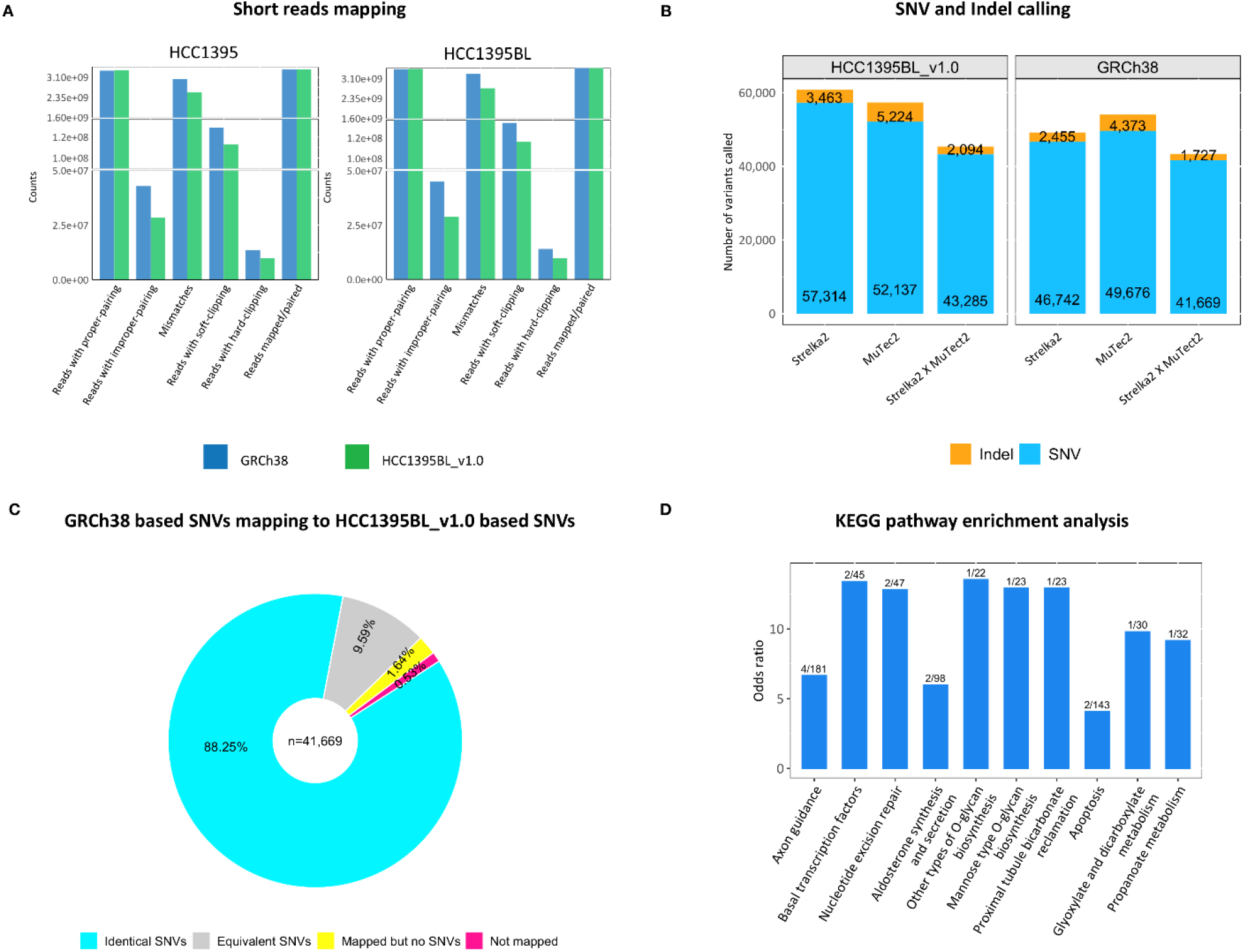
(A) The mappability of short reads on GRCh38 and HCC1395BL_v1.0 with gapped y-axis for visualizing the low range. The numbers of improper-paired reads, reads with mismatches, soft-clipping and hard-clipping were reduced substantially on HCC1395BL_v1.0 for both cell lines. (B) The sensitivity of somatic SNVs and indels detection on GRCh38 and HCC1395BL_v1.0. The numbers of overlapping variants between MuTect2 and Strelka2 were higher on HCC1395BL_v1.0. (C) 41,669 GRCh38 based somatic SNVs mapped onto HCC1395BL_v1.0 based SNVs with 40,768 (97.83%) SNVs as identical (36,773 SNVs, 88.25%) or equivalent (3,995 SNVs, 9.59%). 682 SNVs (1.64%) were able to map on HCC1395BL_v1.0 but without overlapping Strelka2/MuTect2 calls. 219 SNVs (0.53%) were considered as “not-mapped” on HCC1395BL_v1.0 due to the stringent mapping criteria. (D) KEGG pathway enrichment analysis of 71 genes overlapped with the 1,017 novel SNVs detected on HCC1395BL_v1.0 as a reference. Shown here are the top 10 enriched pathways with “Odd Ratio” (zScore) on y-axis. The numeric labels are the enriched gene counts verse the total genes in each pathway.

Based on a previous study [22] and recent SEQC2 reports [46–47], two commonly used somatic mutation callers, namely Strelka2 [23] and MuTect2 [24], were selected for generating reports of somatic SNVs and small indels with the same settings based on same set of Illumina short-read data using HCC1395BL_v1.0 and GRCh38 (alt_loci excluded) as reference genomes, respectively. A higher number of overlapping calls between Strelka2 and MuTect2 were observed, and 1,983 more somatic SNVs/indels were seen on HCC1395BL_v1.0 than GRCh38 (Figure 4B and Supplementary Table 6).

For those 41,669 GRCh38-based somatic SNVs supported by both Strelka2 and MuTect2 callers (Supplementary Table 6), 40,768 SNVs (97.83%) were successfully mapped on HCC1395BL_v1.0 with overlapping SNVs called by Strelka2/MuTect2 (Figure 4C and Supplementary Table 7). 682 SNVs (1.64%) were able to map on HCC1395BL_v1.0 but without overlapping Strelka2/MuTect2 calls. Variant function analysis using ANNOVAR [25] showed that 120 of 682 SNVs were located in exonic or intronic regions, therefore including or excluding these sites would have an impact on downstream mutation interpretations (Supplementary Table 7). 219 SNVs (0.53%) were considered as “not-mapped” on HCC1395BL_v1.0 due to the stringent mapping criteria (see “Methods”). Moreover, 3,995 SNVs (9.58%) were considered as equivalent SNVs between GRCh38 and HCC1395BL_v1.0 but with mismatches in their flanking sequences (Supplementary Figure 4> and Supplementary Table 7). For example, the same set of reads was found (with mapping quality 60) to align across the corresponding intergenic SNV regions on HCC1395BL_v1.0 (scaffold_2:131886469-131886569 for SNV scaffold_2:131886519), but on GRCh38 (chr1:177753949-177754049 for SNV chr1:177753999) two mismatches (or two potentially homozygous germline SNVs) are observed in flanking sequences (Supplementary Figure 4A/4B/4G). Similar examples were found for an exonic SNV at scaffold_37: 17305121 or chr19:17555816 (Supplementary Figure 4C/4D) and an intronic SNV at scaffold_12:48083060 or chr10:114357477 (Supplementary Figure 4E/4F). Such discrepancies reflect the underlined genomic sequence differences between the personalized HCC1395BL_v1.0 and the common reference GRCh38, illustrating the importance of using personal genome for accurate somatic mutation discovery and subsequent analysis such as SNP genotyping assay design and validation as the mismatches in allele-specific probes or primers would impact melting temperature and binding efficiency for a genotyping assay.

Among those 43,285 somatic SNVs supported by both Strelka2 and MuTect2 on HCC1395BL_v1.0 (Supplementary Table 6), 2,790 SNVs were identified without the equivalent GRCh38-based SNVs, and from them, 1,017 sites were well-supported by more than 10 alternate allele reads with the percentage of alternate allele read coverage at least 50%. By co-locating these SNVs with RefSeq genes and transcripts mapped on HCC1395BL_v1.0, 522 of 2,790 SNVs were found in exonic or intronic regions, while 177 SNVs of 1,017 SNVs were overlapping with 71 gene regions (exonic or intronic), suggesting that some meaningful somatic SNVs could be overlooked when using GRCh38 as the reference genome (Figure 4D).

With respect to structural variation identification, more balanced counts of large insertions and deletions were observed on HCC1395BL_v1.0 as reference when alignment-based SV calling tools, such as pbsv (Figure 5A/5B) and Sniffles (Figure 5C/5D) were used. However, as previously reported [14, 26], more insertion calls were identified than deletions on the GRCh38 reference. Such excessive insertion calls were likely related to deletion bias in GRCh38 reference [14, 26].

**Figure 5.**
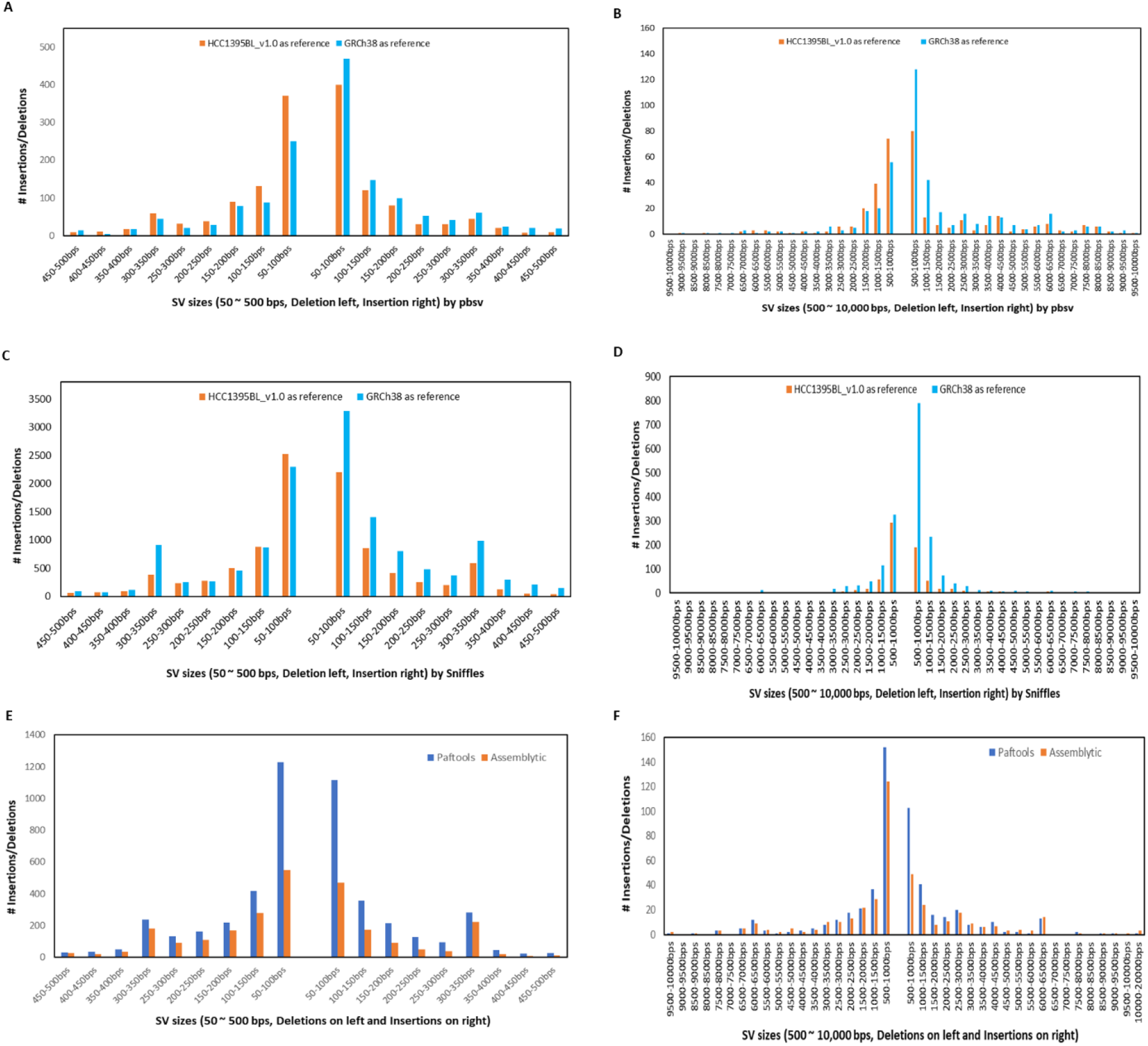
Size distributions of insertions and deletions called by pbmm2/pbsv and Sniffles using HCC1395BL_v1.0 and GRCh38 as references, respectively: (A) 50 – 500 bps by pbsv; (B) 500 – 10,000 bps by pbsv; (C) 50 – 500 bps by Sniffles; (D) 500 – 10,000 bps by Sniffles. Size distribution of SVs detected from direct assembly comparison of HCC1395 to HCC1395BL using paftools and Assemblytics: (E) 50 – 500 bps; (F) 500 – 10,000 bps.

Previously, without a personal genome available, a common reference, such as GRCh38, was the only resource available for assembly-to-assembly based (indirect) structural variation discovery [27]. With an established personal genome based on the normal sample from the same individual, we were able to make a direct comparison of the assembly of the HCC1395 cancer cell line with HCC1395BL_v1.0. We continuously observed more balanced numbers of structural variations (large insertions and deletions), especially for the SVs within the ranges between 50 to 500 bps in this analysis. An *Alu* repeat peak (~300 bps) and an *L1* repeat peak (~6,000 bps) were seen for both insertion and deletion calls (Figure 5E/5F). By merging two alignment-based callsets with two assembly-based callsets and requiring at least two methods supporting each SV site, we identified 3,498 large deletions, 2,239 large insertions and 101 Duplications (DUP) as somatic SVs from HCC1395 cancer cell line.

### Complete mitochondrial genome assemblies reveal non-synonymous somatic mutations

Mitochondria are considered the powerhouse of the cell. However, mitochondrial genome assemblies in recent published *de novo* assemblies have either been absent or highly fragmented [28]. In this study, we completely assembled the mitochondrial genomes for both HCC1395BL and HCC1395 cell lines into single contigs. By directly comparing the two mitochondrial genomes, we identified two undocumented non-synonymous somatic mutations (T4813C and C4938A, not collected in dbSNP153 release) in the MT-ND2 gene, and one non-synonymous somatic mutation (G14249A) on MT-ND6 gene (Figure 6). We manually checked those 3 mutations on the visualization tool IGV and confirmed that those 3 mutations have strong evidence supports. To our knowledge, those 3 somatic mutations were not reported previously in breast cancer [29–32]. Both MT-ND2 and MT-ND6 genes are subunits of the respiratory chain Complex I in the mitochondrial inner membrane, which often contains tumorigenesis related variants [33, 34]; therefore these amino-acid changing alterations may be detrimental to the normal function of the cellular system and should be further investigated.

**Figure 6.**
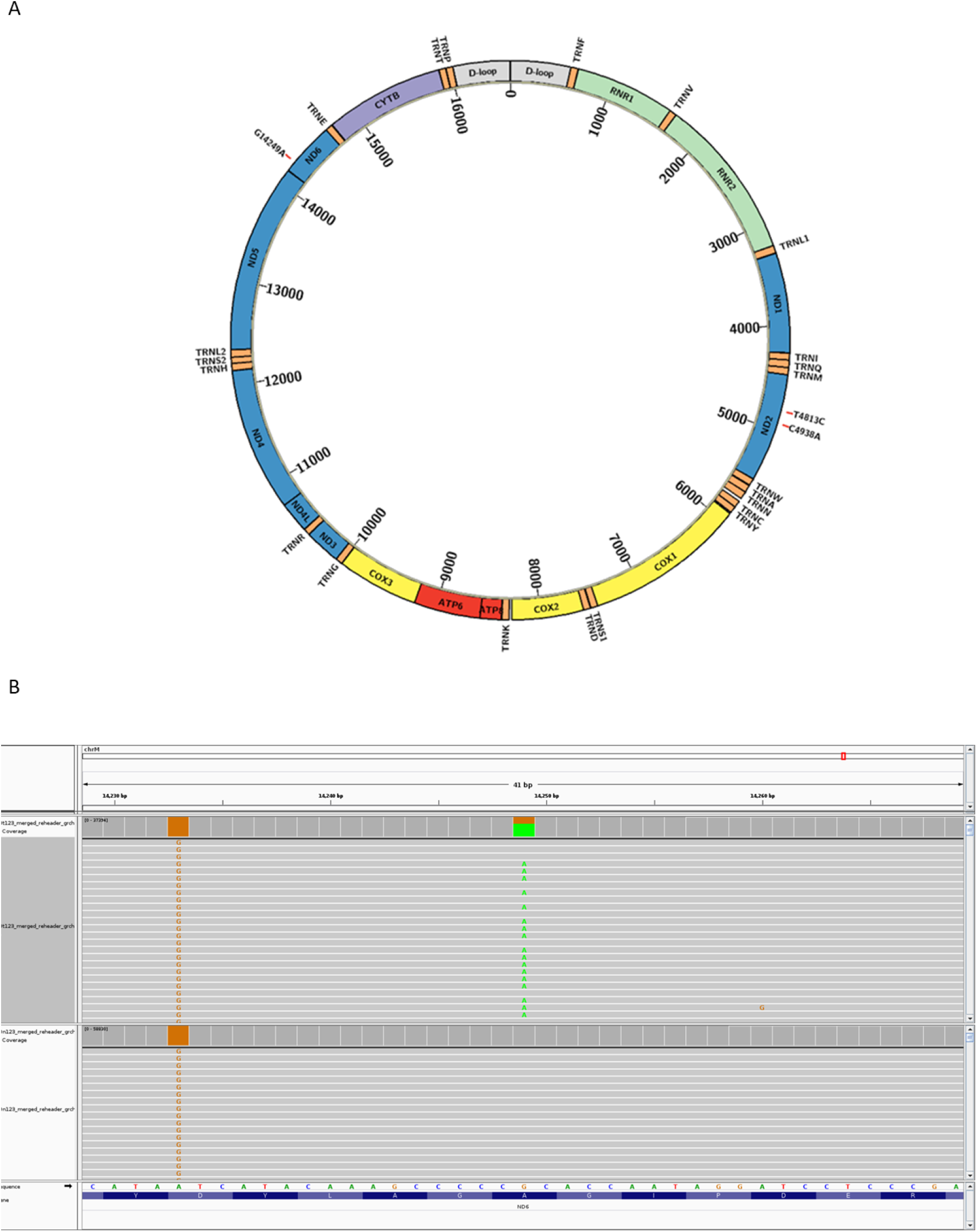
(A) Mitochondrial genome of HCC1395BL/HCC1395 with three unreported non-synonymous somatic mutations (T4813C and C4938A on MT-ND2 gene, and G14249A on MT-ND6 gene), which were confirmed on IGV for G14249A in HCC1395 cell line only, but not in HCC1395BL cell line (B).

## Discussion

In this study, we used a combination of multiple sequencing technologies, including short reads, linked-reads, and long reads to construct the first *de novo* assemblies of a tumor-normal pair from the same individual with breast cancer. We subsequently applied the well-assembled genome as a reference, in comparison to using the common human reference GRCh38, for somatic variation detection and demonstrated the advantages of using a personalized genome as a reference.

Based on our analysis, high quality assembly has been achieved based on our existing data for HCC1395BL in terms of contiguity and gene content. Complex genomic regions have also been well assembled, as we demonstrated that the complete HLA region was assembled into a single scaffold for HCC1395BL with an individualized haplotype. Some clinically relevant genes such as GSTT1 and KIR2DL5 (KIR2DL5A), which are not present in the chromosomes of the GRCh38 primary assembly commonly used for alignment-based analyses, were also captured in our *de novo* HCC1395BL assembly.

For the first time to our knowledge, we were able to identify cancer somatic mutations based on the *de novo* assembly from the same person with breast cancer, instead of inferring them from alignments to a common reference benchmark such as GRCh38 [10]. Our analysis showed that the *de novo* assembly improved read mapping, resulting in a greater percentage of properly-mapped mate pair reads, reducing total numbers of mismatches, and many fewer reads with improper-pairing, soft-clipping, and hard-clipping. Discovery of somatic SNVs and small indels was also more consistent by different calling algorithms, and large insertions/deletions were more balanced in numbers when the *de novo* personal assembly was used for variant calling from this paired reference samples. Mapping analysis of GRCh38-based somatic SNVs set with flankings to the *de novo* assembly revealed that small percentage (1.64%) of GRCh38 based SNVs have good mapping locations on personal genome but without corresponding SNV calls, indicating potential false positives exist. Some SNVs (9.59%) may have the same reference/alternate alleles on *de novo* assembly as on GRCh38 reference genome, but their flanking sequences may be slightly different, highlighting the critical needs of promoting the personal genome assembly for individualized medical research.

Furthermore, we also finished the first complete mitochondrial genomes for a tumor-normal pair. As a result, we were able to perform genome-wide comparative analysis of these two genomes on the personal genome level, resulting in more accurate and complete results from nuclear genome to mitochondrial genome. Our results illustrate that using the *de novo* genome for somatic variant discovery from tumor-normal paired data is possible. We showed that accurate detection of mutation enabling precision cancer medicine can be achieved with a personalized genome, though the overall cost of the sequencing and subsequent computational analysis is substantial.

As sequencing technology continues to evolve with longer read length and lower per-base error rate, many genomic regions such as telomeres and centromeres that were previously impossible to assemble, and whose biology is consequently poorly understood, are now within reach of being finished [14]. In fact, Human Pangenomes Consortium (https://humanpangenome.org/) has been working on constructing a more representative genome reference, with the goal of producing 300 high quality haplotype resolved human genomes from different human sub-populations. Such new developments should encourage the Consortium and other researchers to continue improving the quality of the *de novo* personal assembly for this tumor-normal pair by applying PacBio’s HiFi reads and Oxford Nanopore’s ultra-long reads in the near future. Such advancement in genome assembly will offer a better path forward for somatic variant identification using personalized genome as a reference, and thus provide more insights into understanding how tumorigenesis occurs from the molecular level, leading to discovery of vital genetic markers for cancer diagnostic and drug development.

## Methods

### Whole genome sequencing datasets

A matched tumor/normal pair of TNBC breast cancer cell lines (HCC1395 as tumor and HCC1395BL as normal) was selected as reference samples for whole genome sequencing with multiple platforms. For HCC1395BL assembly analysis, included were about 175-fold of Illumina short reads, 161-fold of 10X Genomics (10X) linked reads, 53-fold of Pacific Bioscience (PacBio) long reads, 71-fold of Hi-C reads, and 15-fold Oxford Nanopore technologies (ONT) reads, respectively (Table S1). For HCC1395, about 170-fold of Illumina short reads, 160-fold of 10X Genomics linked reads, and 46-fold of Pacific Bioscience (PacBio) long reads were included. Library preparations and sequencing for Illumina short reads, 10X linked reads and PacBio long reads were described previously [46–47].

Dovetail HiC library preparation and sequencing: The Dovetail HiC libraries were prepared for each sample in a similar manner as described previously (Erez Lieberman-Aiden et al., 2009). For each library, chromatin was fixed in place with a 1% formaldehyde solution in the nucleus and then extracted. Fixed chromatin was digested with DpnII, the 5’ overhangs were filled in with biotinylated nucleotides, and then free blunt ends were ligated. After ligation, crosslinks were reversed, and the DNA purified from protein. Purified DNA containing biotinylated free-ends were removed as those are not reflective of proximity-ligated molecules. The DNA was then sheared to ~350 bp mean fragment size and sequencing libraries were generated using NEBNext Ultra enzymes and Illumina-compatible adapters. Internal biotin-containing fragments were isolated using streptavidin beads before PCR enrichment of each library. The libraries were sequenced on an Illumina HiSeq X to a depth of ~200M read pairs per library.

Oxford Nanopore technologies (ONT) MinION sequencing data: Genomic DNA from HCC1395BL cell line was extracted using QIAGEN MagAttract HMW DNA Kit (QIAGEN, Hilden, Germany). 1ug of initial DNA without fragmentation was used for library construction using SQK-LSK109 ligation sequencing kit (Oxford Nanopore Technologies, Oxford, UK). Library preparations were conducted as per the protocols provided by ONT. Each library was sequenced on an individual MinION FLO-MIN106D R9.4 flowcell. Prior to sequencing, flowcell pore counts were measured using the MinKNOW Platform QC script (Oxford Nanopore Technologies, Oxford, UK). About 300ng of completed libraries were loaded as per instructions from ONT. Raw sequence reads were basecalled in real time via MinKNOW. Basecalled data passing quality parameters (qmean > 7) were converted to fastq. Only reads designated as pass were included in further analyses.

### Assembly, polishing, Scaffolding, and Phasing

PacBio long reads data was firstly error-corrected and then assembled into primary contigs using “canu” assembler (version 1.8) [17]. The contig sequences were then polished with Illumina paired-end reads using PILON (version 1.22) [35]. The polishing process was performed twice to achieve the best results. Linked reads from 10X Genomics were assembled using “Supernova” assembler (version 2.0.0) [18]. Scaffolding with linked-reads was performed using ARCS (version 1.0.5) [36], while scaffolding with Hi-C data was completed using SALSA (https://github.com/marbl/SALSA) [37].

After scaffolding with ARCS and SALSA, we mapped the unitig sequences, which were produced with Illumina short reads using fermikit (version r188) [38], to the scaffold assembly using BWA [39], and then used bcftools (version 1.6, https://samtools.github.io/bcftools/bcftools.html) to generate the final consensus assembly.

The Illumina short reads were aligned onto the HCC1395BL_v1.0 using BWA mem [39], and duplicated reads were marked with Picard MarkDuplicates. High confident heterozygous sites were identified using GATK4 (version gatk-4.0.3.0) [40]. Phasing was performed with the identified high confident heterozygous sites and long reads from PacBio and ONT using phasing tools WhatsHap (version 0.18) [41]. Two haplotypes of the assembly in FASTA format were also reconstructed with the phasing information. Assembly-based germline SNVs in autosomal chromosomes were called using dipcall (https://github.com/lh3/dipcall) with two haplotypes as inputs.

### Assembly evaluation

QUAST (version 5.0.0) [42], Benchmarking Universal Single-Copy Orthologue (BUSCO, version 3.0.0) [43] were used to assess the quality of each de novo assembly. BLAT (v36) was used for mapping all RefSeq mRNA transcripts (accession prefixed with NM_ and NR_) (ftp://ftp.ncbi.nlm.nih.gov/genomes/all/GCF/000/001/405/GCF_000001405.38_GRCh38.p12/GCF_000001405.38_GRCh38.p12_rna.fna.gz) that were previously annotated on the GRCh38 assembly to the new assembly with parameter minIdentity 92.

For GRCh38 consistency analysis, each assembly was compared with GRCh38 reference assembly (ftp://ftp-trace.ncbi.nlm.nih.gov/ReferenceSamples/seqc/technical/reference_genome/GRCh38/GRCh38.d1.vd1.fa) using minmap2 [44]. Alignments with mapping quality 60 and alignment length 100Kb+ were considered as good links for the consistency plot by Circos (Krzywinski, M., et al., 2009). Scaffolds smaller than 10 kb were excluded from the analysis.

We also introduced a new parameter “Top50”, which is the summed length of the 50 longest scaffolds, to monitor the contiguity of a given assembly during scaffolding process, as the long-read and Hi-C sequencing technologies could make it possible to have arm-scale or chromosomal scale assembly. For the human genome with a total of 48 chromosomal arms, Top50 might be a suitable indicator to reflect the contiguity of the scaffold assembly if each chromosomal arm forms a scaffold.

### Genome annotation

To better annotate the final assembly HCC1395BL_v1.0, BLAT (version 36) and AUGUSTUS (version 3.3.1) pipeline [45] was used to map the previously described RefSeq transcripts to the assembly (excluding all pseudogenes and genes from NC_000024 chromosome Y). Protein-coding transcripts with annotations containing “pseudogene” and non-protein-coding transcripts with annotations containing “pseudo=true” in their deflines were considered as “pseudogenes” in this analysis. For BLAT, the option “minIdentity” was set to 92. Transcripts with more than 95% alignment coverage and 95% ungapped identity were considered mapped on HCC1395BL_v1.0 assembly. One exception was applied to HLA-DQA1 (NM_002122.3) gene that was 100% covered by the HCC1395BL_v1.0, but with the identity of 92.95%, for comparing the HLA haplotypes between GRCh38 and HCC1395BL_v1.0. No other mapped location was found for HLA-DQA1 gene on HCC1395BL_v1.0. In case of multiple mapping locations, the best mapping location with maximum number of matching bases for the transcript was selected.

### Somatic variant detection

BWA [39] was used to align the Illumina short reads onto the *de novo* assembly (HCC1395BL_v1.0) and GRCh38 primary assembly, respectively. Duplicate reads were marked with Picard MarkDuplicates. Mapping statistics was gathered using samtools with the “stats” option (http://www.htslib.org/doc/samtools-stats.html).

Strelka2 (version 2.9.2) [23] and MuTect2 (version gatk-4.0.3.0/gatk Mutect2) [24] were used to generate somatic SNVs and indels. The chrX, chr6p and chr16q regions were not included for somatic variant comparison, for consistency with the reference somatic set (v1.1) from the SEQC2 Somatic Mutation Working Group [46–47].

Structural variations from PacBio long reads data were identified using pbmm2/pbsv (version 2.2.1, https://github.com/PacificBiosciences/pbsv) and Sniffles (version 1.0.11, https://github.com/fritzsedlazeck/Sniffles). For each reference assembly, SVs were called separately using PacBio long read alignments of HCC1395 and HCC1395BL, then the SV calls of HCC1395 were filtered with SVs of HCC1395BL. Sniffles callsets were filtered with AF >= 0.9 and 10 ~ 100 read supports.

Assembly-based SVs were generated from direct assembly comparisons of HCC1395 cancer cell line contig assembly with the *de novo* HCC1395BL assembly using paftools [44] and Assemblytics (https://github.com/MariaNattestad/assemblytics). For consistency with SEQC2 Somatic Mutation Working Group [46–47], calls from chrX, chr6p, and chr16q were also excluded for comparison. SURVIVOR (version 1.0.7, https://github.com/fritzsedlazeck/SURVIVOR) was used for merging and filtering SV callsets.

### Mapping GRCh38 SNVs to *de novo* assembly

Two-steps mapping approaches were performed to find corresponding locations of the GRCh38-based somatic SNVs on the *de novo* assembly. We extracted both the reference and alternate alleles of each SNV with their 50bps flanking sequences from GRCh38 and created a fasta file before mapping using BLAST (blast 2.10.1). The first step was to map all SNVs with higher criteria so that SNVs with identity >=99% and alignment length >=101 bps were selected. The un-selected SNVs from Step1 were then mapped in the second step with lower thresholds (using 95% identity and 95 bps alignment coverage as cut-off based on our mapping experiments) so that best mapped SNVs with some mismatches and small indels would be selected (Figure S6). For both steps, both alleles of each SNV were required to map onto the same locations with identical start and end on *de novo* assembly. In addition, to be considered as an equivalent SNV call between GRCh38 and *de novo* assembly, the alternate allele must be at the center position. Manual inspections on IGV for some SNVs were also performed. Unselected SNVs from Step2 were considered as unmapped.

### Variant annotation and pathway analysis

Variant function analysis was performed using ANNOVAR [25]. Pathway analysis were performed through Enrichr web site (https://maayanlab.cloud/Enrichr/).

### Mitochondrial sequence analysis

Contigs from HCC1395BL assembly and HCC1395 assembly that were fully covering the mitochondrial sequences from GRCh38 (16,569 bps; https://www.ncbi.nlm.nih.gov/nuccore/NC_012920.1) were selected based on minimap2 mapping results. Since the mitochondrial genome is circular, the full mitochondrial sequences were extracted from each of the selected contigs based on BLAST mapping results. CLUSTAL (v1.2.4) was used to generate multiple sequence alignments for variant analysis. The variants were annotated with the MITOMAP human mitochondrial genome database (http://www.mitomap.org,2019) and dbSNP (v153).

## Supporting information

Supplementary_Materials_April08_2021

## Data availability

All raw data are available in NCBI SRA database (SRP162370). The HCC1395BL_v1.0 assembly has been deposited at DDBJ/ENA/GenBank under the accession JAEOAY000000000. Final *de novo* assembly fasta file and *de novo* assembly-based somatic mutation VCF files are also accessible via the NCBI ftp site (https://ftp://ftp-trace.ncbi.nlm.nih.gov/ReferenceSamples/seqc/Somatic_Mutation_WG/assembly).

## Code and software availability

All software or tools used for *de novo* genome assemblies, assembly evaluations, and variant calls were publically available and listed in “Methods” section.

## Acknowledgements

The genomic work carried out at the LLU Center for Genomics was funded in part by the National Institutes of Health (NIH) grant S10OD019960 (CW). The study was also partially supported by the American Heart Association grant 18IPA34170301 (CW). Chunlin Xiao and Valerie Schneider were supported by the Intramural Research Program of the National Library of Medicine, National Institutes of Health. Javkhlan Ganbat provided useful help regarding HiC data analysis.

## Author contributions

CX, WX designed the study and wrote the manuscript draft. CP performed HiC sequencing and QA/QC. ZC, WC, TL and CW performed ONT sequencing and QA/QC. CX, WX, WC, ZC, CW, and LF performed the analyses. CX, WX, WC, ZC, CW, CP, and VS improved the manuscript. CX, VS and WX managed the data.

## Disclaimer

The content of this manuscript is solely the responsibility of the authors, and the views presented here do not necessarily reflect official policy of the US Food and Drug Administration or US National Institutes of Health. Any mention of commercial products or materials or tools is purely for clarification purpose and not intended as endorsement or discouragement.

