## Supplementary_Materials_April08_2021 for "Personalized genome assembly for accurate cancer somatic mutation discovery using cancer-normal paired reference samples"

### Supplementary Figures

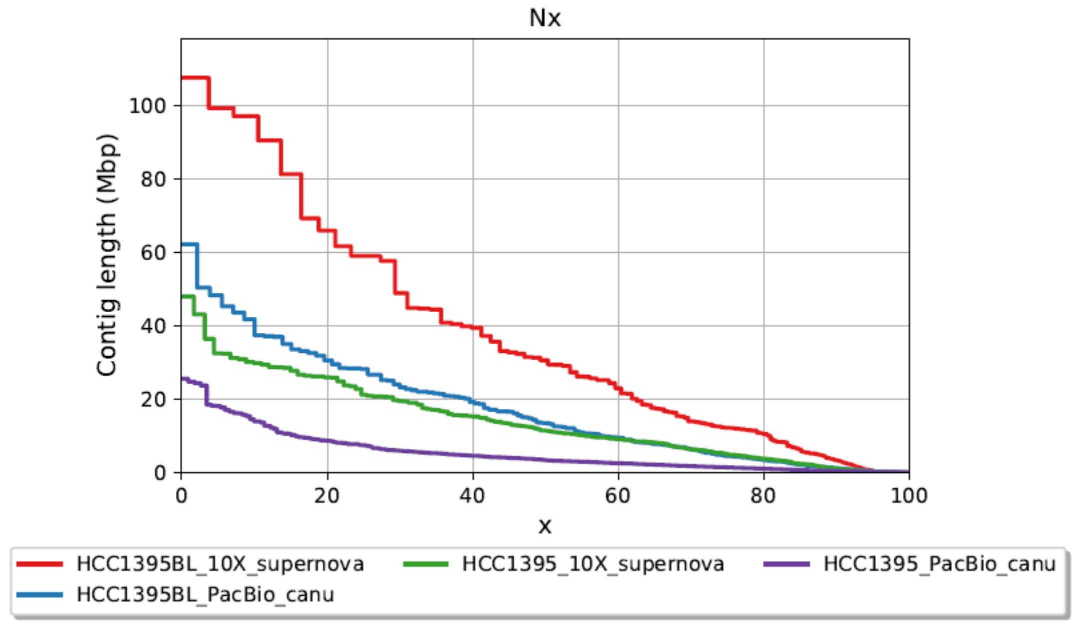

Supplementary Figure 1. Nx plot of the assemblies from PacBio long reads using canu and 10X Genomics linked reads using supernova.

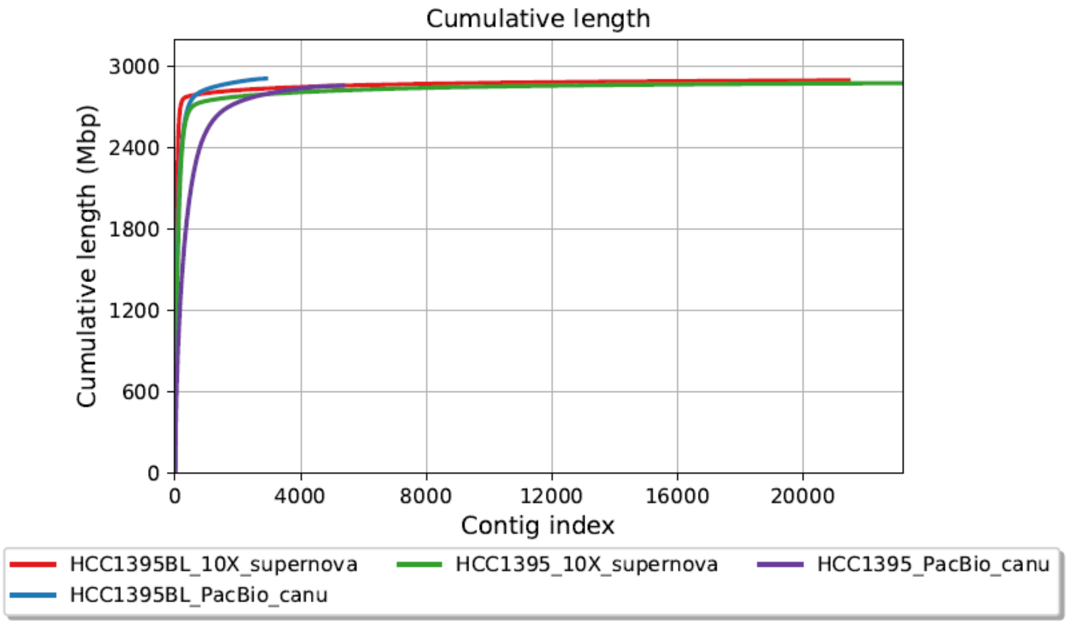

Supplementary Figure 2. Cumulative length plot of the assemblies from PacBio long reads using canu and 10X Genomics linked reads using supernova.

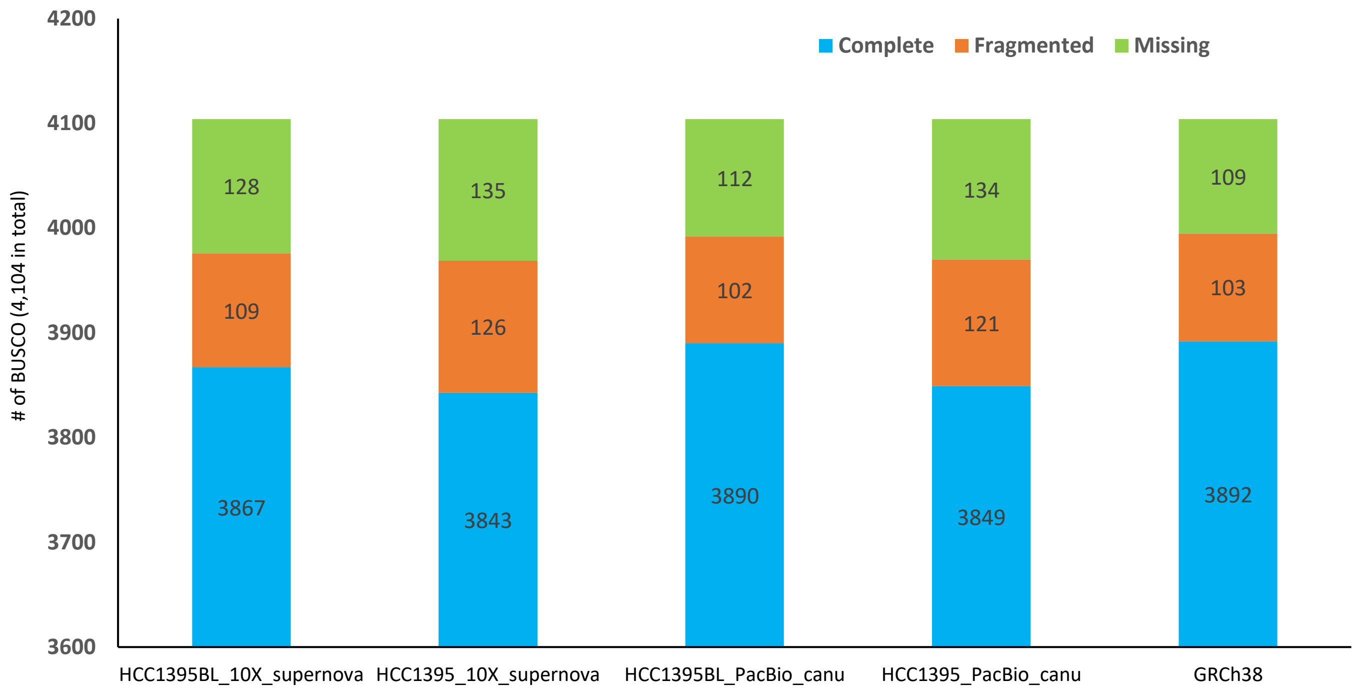

Supplementary Figure 3. BUSCO assessment of the assemblies from PacBio long reads using canu and 10X Genomics linked reads using supernova. GRCh38 (primary assembly) was listed here for comparison purposes.

**A. Intergenic SNV scaffod_2:131886519 on HCC1395BL_v1.0 assembly.**

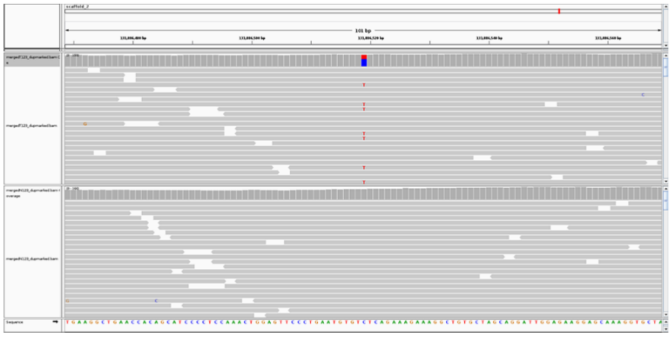

**B. Intergenic SNV chr1:177753999 on GRCh38 with mismatches in flanking.**

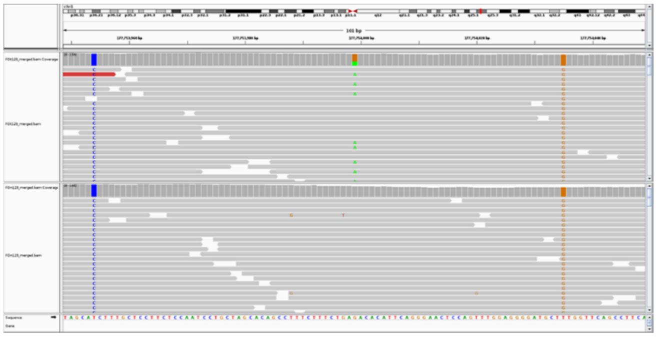

**C. Exonic SNV scaffold_37: 17305121 on HCC1395BL_v1.0 assembly.**

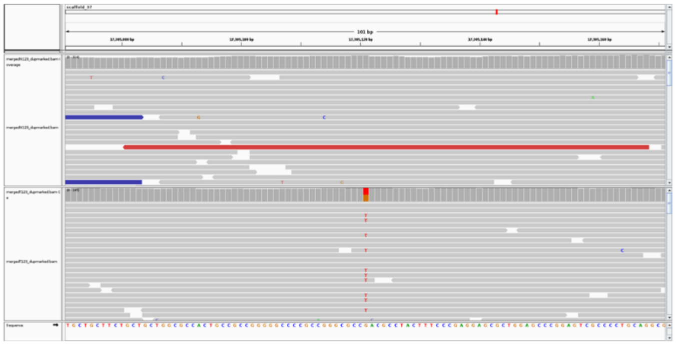

**D. Exonic SNV chr19:17555816 on GRCh38 with mismatches in flanking.**

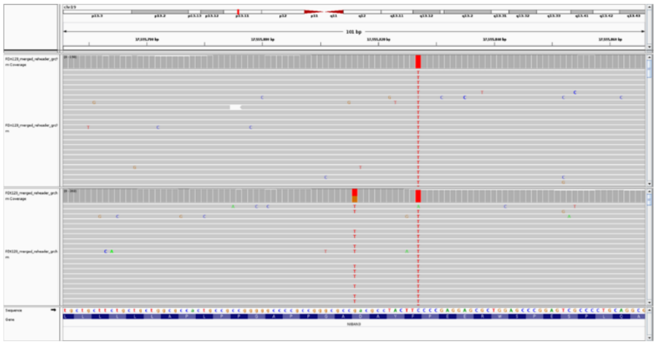

**E. Intronic SNV scaffold_12:48083060 on HCC1395BL_v1.0 assembly.**

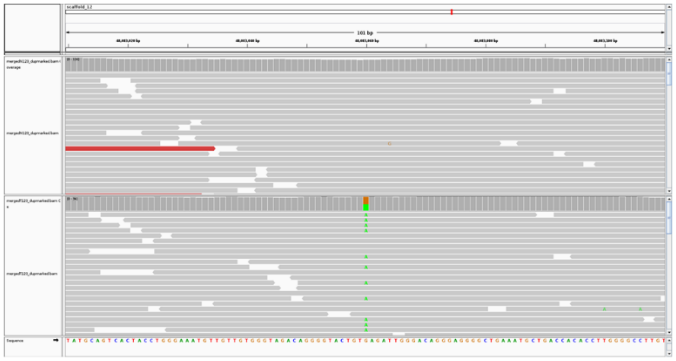

**F. Intronic SNV chr10_114357477 on GRCh38 with mismatches in flanking.**

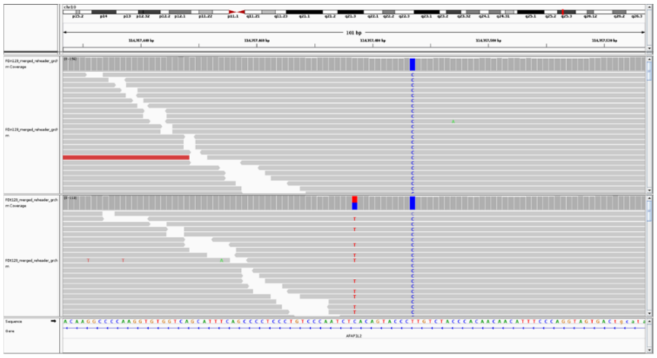

**G.** BLAST alignment example of reference allele and alternate allele from SNV chr1:177753999 mapped onto de novo assembly.

```
Query= chr1_177753999_ref

Length=101
                                                                      Score     E
Sequences producing significant alignments:                          (Bits)  Value

scaffold_2                                                            176     2e-42

>scaffold_2
Length=159912780

 Score = 176 bits (95),  Expect = 2e-42
 Identities = 99/101 (98%), Gaps = 0/101 (0%)
 Strand=Plus/Minus

Query  1          TAGCATCTTTGCTCCTTCTCCAATCCTGCTAGCACAGCCTTTCTTTCTGAGACACATTCA  60
                  ||||| ||||||||||||||||||||||||||||||||||||||||||||||||||||||
Sbjct  131886569  TAGCACCTTTGCTCCTTCTCCAATCCTGCTAGCACAGCCTTTCTTTCTGAGACACATTCA  131886510

Query  61         GGGAACTCCAGTTTGGAGGGGATGCTTTGGTTCAGCCTTCA  101
                  |||||||||||||||||||||||||| ||||||||||||||
Sbjct  131886509  GGGAACTCCAGTTTGGAGGGGATGCTGTGGTTCAGCCTTCA  131886469

Query= chr1_177753999_alt

Length=101
                                                                      Score     E
Sequences producing significant alignments:                          (Bits)  Value

scaffold_2                                                            171     7e-41

>scaffold_2
Length=159912780

 Score = 171 bits (92),  Expect = 7e-41
 Identities = 98/101 (97%), Gaps = 0/101 (0%)
 Strand=Plus/Minus

Query  1          TAGCATCTTTGCTCCTTCTCCAATCCTGCTAGCACAGCCTTTCTTTCTGAAACACATTCA  60
                  ||||| |||||||||||||||||||||||||||||||||||||||||||| |||||||||
Sbjct  131886569  TAGCACCTTTGCTCCTTCTCCAATCCTGCTAGCACAGCCTTTCTTTCTGAGACACATTCA  131886510

Query  61         GGGAACTCCAGTTTGGAGGGGATGCTTTGGTTCAGCCTTCA  101
                  |||||||||||||||||||||||||| ||||||||||||||
Sbjct  131886509  GGGAACTCCAGTTTGGAGGGGATGCTGTGGTTCAGCCTTCA  131886469
```

Supplementary Figure 4.  Illustration of GRCh38 SNV mapping to *de novo* assembly with discrepancies.

(A) IGV snapshot for an intergenic SNV scaffold_2:131886519, the alleles were reverse-complement mapped on *de novo* assembly.

(B) IGV snapshot for SNV chr1:177753999 on GRCh38 with mismatches in flanking sequences. The same set of reads (266 reads from HCC1395BL sample, and 278 reads from HCC1395 sample) were found to align (with mapping quality 60) crossing the corresponding SNV regions (scaffold_2:131886469-131886569 for SNV scaffold_2:131886519 on HCC1395BL_v1.0; chr1:177753949-177754049 for SNV chr1:177753999 on GRCh38).

(C) IGV snapshot for an exonic SNV scaffold_37: 17305121 on HCC1395BL_v1.0 assembly.

(D) IGV snapshot for an exonic SNV chr19:17555816 on GRCh38 with mismatches in flanking. This somatic SNV causes amino acid change (Asp -> Tyr for GAC -> TAC) in gene COLGALT1 on chr19.

(E) IGV snapshot for an intronic SNV scaffold_12:48083060 on HCC1395BL_v1.0 assembly; (F) IGV snapshot for an intronic SNV chr10_114357477 on GRCh38 with mismatches in flanking. This somatic SNV is located in the intronic region of the gene AFAP1L2 on chr10.

(G) BLAST alignment example of reference allele and alternate allele from SNV chr1:177753999 mapped onto de novo assembly with reference allele 98% identity and 101 bps alignment for reference allele, and 97% identity and 101 bps alignment for alternate allele.

### Supplementary Tables

Supplementary Table 1. Whole genome sequencing data of HCC1395 and HCC1395BL by multiple sequencing platforms used in this study.

| Dataset | Sequencing center | Usage | HCC1395 | | HCC1395BL | |
|---|---|---|---|---|---|---|
| | | | # Reads | Total base pairs (Depths) | # Reads | Total base pairs (Depths) |
| Illumina HiSeq [ref 46-47] | Fudan (FD) | Polishing, unitig assembly, variant calling | 3,403,236,080 | 510,485,412,000 (170.16x) | 3,511,999,736 | 526,799,960,400 (175.59x) |
| 10X Genomics Chromium [ref 46-47] | Fudan (FD) | Assembly; Scaffolding | 3,206,110,814 | 480,916,622,100 (160.30x) | 3,222,458,614 | 483,368,792,100 (161.12x) |
| PacBio Sequel [ref 46-47] | CSHL | Primary Contig assembly, phasing, variant calling | 17,265,341 | 140,649,790,223 (46.88x) | 17,671,082 | 160,626,774,754 (53.54x) |
| Hi-C | Dovetail | Scaffolding | NA | NA | 1,432,923,164 | 214,938,474,600 (71.64x) |
| Oxford Nanopore | Loma Linda University | Phasing | NA | NA | 11,585,856 | 47,006,385,877 (15.66x) |

Supplementary Table 2: Summary of the *de novo* assemblies for the two cell lines using PacBio long reads and 10X Genomics linked reads.

| | **HCC1395BL (10X_supernova)** | **HCC1395 (10X_supernova)** | **HCC1395BL (PacBio_canu)** | **HCC1395 (PacBio_canu)** |
|---|---|---|---|---|
| # contigs (>= 0 bp) | 21,450 | 23,195 | 2,900 | 5,351 |
| Total length (>= 0 bp) | 2,893,054,450 | 2,871,887,885 | 2,905,219,353 | 2,853,495,799 |
| # contigs (>= 10,000 bp) | 3,416 | 4,666 | 2,828 | 4,949 |
| Total length (>= 10,000 bp) | 2,837,767,582 | 2,812,618,703 | 2,904,842,414 | 2,851,315,749 |
| Largest contig | 107,564,373 | 47,962,721 | 62,208,403 | 25,523,694 |
| GC (%) | 40.89 | 40.96 | 40.91 | 40.95 |
| N50 | 30,593,498 | 11,455,133 | 13,480,407 | 3,338,913 |
| L50 | 27 | 70 | 57 | 210 |
| # contigs (>= 10,000 bp) matched GRCh38 | 3,176 | 4,449 | 2,526 | 4,627 |
| # bps of contig (>= 10,000 bp) matched GRCh38 | 2,833,045,730 | 2,808,106,025 | 2,888,438,712 | 2,836,167,093 |
| # cumulative bps matches on GRCh38 | 2,787,709,216 | 2,754,283,911 | 2,867,274,440 | 2,820,906,016 |
| # novel contigs (>= 10,000 bp) | 240 | 217 | 302 | 322 |

| | | | | |
|---|---|---|---|---|
| # bps of novel contigs (>= 10,000 bp) | 4,721,852 | 4,512,678 | 16,403,702 | 15,148,656 |
| RefSeq NMs mapped (95+% alignment covered) | 48,061 (96.02%) | 47,128 (94.15%) | 49,287 (98.41%) | 47,619 (95.13%) |
| RefSeq NRs mapped (95+% alignment covered) | 14,423 (92.78%) | 14,165 (91.12%) | 15,115 (97.24%) | 14,717 (94.67%) |

Supplementary Table 3. The summary of genome phasing using WhatsHap with PacBio reads alone and reads from combinations of PacBio and Oxford Nanopore (ONT).

| | PacBio reads | PacBio + ONT reads |
|---|---|---|
| # Blocks | 6,368 | 3,204 |
| Sum block length (bps) | 2,421,024,137 | 2,543,961,355 |
| Longest block (bps) | 6,378,479 | 20,455,447 |
| # Phased | 3,134,631 | 3,135,816 |
| # Hets in VCF | 3,172,233 | 3,172,233 |

Supplementary Table 4. Summary of RefSeq genes and transcripts mapping onto HCC1395BL_v1.0 assembly. Genes from ChrY and all pseudogenes were excluded from this analysis. For comparison, the same set of RefSeq sequences were mapped onto GRCh38.

| | # RefSeq NM | # genes for RefSeq NM | # RefSeq NR | # genes for RefSeq NR |
|---|---|---|---|---|
| # Input (excluding chrY, and all pseudo genes) | 49,844 | 19,325 | 14,132 | 10,061 |
| # Found on HCC1395BL_v1.0 | 49,838 | 19,322 | 14,131 | 10,060 |
| # Not Found on HCC1395BL_v1.0 | 6 | 3 | 1 | 1 |
| # Found 50%+ coverage & 95%+ identity on HCC1395BL_v1.0 | 49,798 (99.91%) | 19,303 (99.89%) | 14,120 (99.92%) | 10,049 (99.88%) |
| # Found 95%+ coverage & 95%+ identity on HCC1395BL_v1.0 | 49,482 (99.27%) | 19,164 (99.17%) | 13,978 (98.91%) | 9,958 (98.98%) |
| # Found 50%+ coverage & 95%+ identity on GRCh38 | 49,825 (99.96%) | 19,318 (99.96%) | 14,128 (99.97%) | 10,058 (99.97%) |
| # Found 95%+ coverage & 95%+ identity on GRCh38 | 49,669 (99.65%) | 19,232 (99.52%) | 14,044 (99.37%) | 10,002 (99.41%) |

Supplementary Table 5. Summary of Illumina short read mappings on HCC1395BL_v1.0 assembly as opposed to GRCh38.

| | HCC1395BL cell line | | HCC1395 cell line | |
|---|---|---|---|---|
| | GRCh38 as reference | HCC1395BL_v1.0 as reference | GRCh38 as reference | HCC1395BL_v1.0 as reference |
| # Reads with proper-pairing | 3,453,531,324 | 3,469,571,424 | 3,348,661,428 | 3,362,223,188 |
| # Reads with improper-pairing | 45,094,500 | 28,819,322 | 42,837,478 | 28,403,442 |
| # Mismatches (from BAM NM tag) | 3,287,384,223 | 2,736,108,247 | 3,034,142,098 | 2,545,029,100 |
| # Reads with soft-clipping | 134,583,157 | 116,428,804 | 129,114,705 | 113,160,104 |
| # Reads with hard-clipping | 14,076,941 | 9,704,136 | 13,634,238 | 9,820,328 |
| # Reads mapped/paired | 3,498,625,824 | 3,498,390,746 | 3,391,498,906 | 3,390,626,630 |

Supplementary Table 6. Summary of somatic SNVs/Indels detected by Strelka2 and MuTect2 on GRCh38 and HCC1395BL assembly, respectively.

| | **HCC1395BL_v1.0 as reference** | | | **GRCh38 as reference** | | |
|---|---|---|---|---|---|---|
| | Strelka2 | MuTect2 | Strelka2 / MuTect2 intersection | Strelka2 | MuTect2 | Strelka2 / MuTect2 intersection |
| # SNVs | 57,314 | 52,137 | 43,285 | 46,742 | 49,676 | 41,669 |
| # Indels (1~20 bps) | 3,463 | 5,224 | 2,094 | 2,455 | 4,373 | 1,727 |
| # Total | 60,777 | 57,361 | 45,379 | 49,197 | 54,049 | 43,396 |

Supplementary Table 7. Summary of mapping 41,669 GRCh38-based SNVs (intersection of Strelka2/MuTect2 SNVs) to *de novo* assembly HCC1395BL_v1.0, and their function annotation using ANNOVAR.

| | Mapped on de novo assembly WITH overlapping Strelka2/MuTect2 SNVs | | Mapped on de novo assembly WITHOUT overlapping Strelka2/MuTect2 SNVs | Not-mapped on *de novo* assembly | Total |
|---|---|---|---|---|---|
| | No mismatch in flanking | Mismatch in flanking | | | |
| UTR3 | 363 | 28 | 2 | 1 | 394 |
| UTR5 | 107 | 2 | 1 | 1 | 111 |
| downstream | 323 | 36 | 7 | 2 | 368 |
| exonic | 442 | 28 | 5 | 2 | 477 |
| intergenic | 17698 | 2112 | 480 | 135 | 20425 |
| intronic | 12726 | 1297 | 115 | 56 | 14194 |
| ncRNA_exonic | 264 | 24 | 3 | 2 | 293 |
| ncRNA_intronic | 4479 | 432 | 62 | 19 | 4992 |
| upstream | 360 | 34 | 7 | 1 | 402 |
| splicing | 11 | 2 | 0 | 0 | 13 |
| **Total # SNVs** | **36,773 (88.25%)** | **3,995 (9.59%)** | **682 (1.64%)** | **219 (0.52%)** | **41,669 (100%)** |
